# LLM-Assisted Functional Gene Annotation

**DOI:** 10.1101/2025.08.06.668957

**Authors:** Calvin Boyce, Clara Pereira, Gyeong Dae Kim, Allegra Petti

**Author notes:** Contributing authors.

## Abstract

Functional gene annotation is a highly manual and subjective process that requires analysis of large amounts of statistical and literature results. Here we present Artificial Intelligence Gene Enrichment (AIGE), a careful automation of the current state of the art process used by bioinformatic experts to determine functions enriched in novel or experimentally derived gene lists. In 1206 test cases, AIGE is able to accurately recover 87% of biological functions, pathways, and cell types, is robust to noise contamination, and accurately assesses its own self-confidence. AIGE reports provide accurate functional annotations and encourage further research in a broad set of contexts.

## 1 Introduction

Functional gene enrichment is a fundamental task in genomic and molecular biology, and an instrumental step in most standard bioinformatic analyses. Extensive work has been done to determine the functions and characteristics of individual genes[1, 2], and to compile lists of genes associated with specific biological or molecular functions[3], but elucidating the phenotypic behavior of a novel gene set in the context of a specific biological question requires carefully designed experimentation. Large scale bioinformatic analyses often include dozens or even hundreds of novel gene lists for which individual experimentation to determine functional enrichment is infeasible. In these settings, approximate functional annotations are often obtained by comparing novel gene lists against databases of known gene lists to determine statistical significance of overlap via either Over-Representation Analysis (ORA) or Gene Set Enrichment Analysis (GSEA)[4]. ORA uses Fisher’s exact test or a hypergeometric test to compare the overlap between a set of genes and a database of predefined gene lists, and is often paired with Bonferroni or false discovery rate statistical correction to account for multiple comparisons. GSEA determines the over or underrepresentation of a known set of genes in a ranked list of Differentially Expressed Genes (DEGs) using a Kolmogorov-Smirnov-like running sum. Each of these methods determines the likelihood of observing a degree of overlap between the novel gene set and the known gene lists under the assumption that genes are expressed uniformly and independently from one another, a strong assumption that leaves room for statistical innovation.

Several tools and methods have been built to perform these statistical comparisons including ToppFun[5], Enrichr[6], clusterProfiler[7], g:Profiler[8], GSEA[4], and fgsea[9], providing enriched terms from Gene Ontology (GO)[3], Kyoto Encyclopedia of Genes and Genomes (KEGG)[10], Molecular Signatures Database (MSigDB)[11], and other curated lists of genes and their functional or pathway involvement. While these results give researchers a good statistical starting point, a proper functional enrichment analysis often includes extensive literature review to verify the functions of subsets of genes in the novel list, and an expert opinion to successfully synthesize the available information. This is a highly manual and subjective process that significantly slows down large-scale research and analysis.

With the advent of Large Language Models (LLMs) and their ability to parse textual data at scale, several recent approaches have applied LLMs to bioinformatic tasks such as cell typing[12–16] and functional gene enrichment[17–20]. While the more novel approaches perform fundamental pre-training similar to that of Generative Pre-Trained (GPT) models[13], or integrate literature sources to augment biological knowledge[21], most approaches use flagship foundation models from sources like OpenAI, Anthropic, or DeepSeek out of the box and evaluate their performance reproducing Gene Ontology or other defined gene list names without fine tuning or training (i.e. zero-shot). This strategy takes advantage of the impressive capacity for LLMs to apply domain expertise to novel data in a flexible and scalable format.

The current quintessential objection to using LLMs for zero-shot prediction is the inability to examine the reasons behind the token predictions of transformers, the fundamental unit of modern LLMs, leading to a lag in adoption of AI-based tools in biology and many other fields. While Chain of Thought (CoT) reasoning models provide some insight into the underlying processes of LLMs, recent research has cast skepticism on the reliability of reasoning models to represent their thinking process or generalize to truly novel data[22–24]. In an approach more analogous to classical neuroscience, researchers at Anthropic have recently pioneered an approach to examine “meta-features” (groups of neurons that “fire” or activate together) in their neural networks to better understand the process LLMs use to make decisions, but this field of research is still in its infancy[25].

Most recently, Wang et. al. published GeneAgent[26] which uses zero-shot LLM prediction to propose functions that may be enriched in a set of genes, then passes those proposed functions through an agent to support or refute their validity by comparing them against classic statistical tools and literature. This approach successfully minimizes the hallucinations of the LLM labeling the gene lists, but it also restricts the capacity of the LLM to generate novel annotations not found in existing databases. The validity of a novel claim made by an LLM can only truly be determined experimentally, which is difficult to scale.

Here we propose Artificial Intelligence Gene Enrichment (AIGE), an LLM-augmented automation of the current expert best practice for gene annotation. The model follows the same steps a human scientist would by parsing statistical sources provided by ToppFun, Enrichr, and g:Profiler, as well as gene summaries and literature findings from NCBI’s gene and PubMed databases to produce a holistic report that determines functions enriched in a novel set of genes while providing industry standard citations. This approach only utilizes the LLM for data synthesis, not for claim generation. It does not take advantage of the capacity for LLMs to propose previously unannotated functions, but it also eliminates the capacity for hallucination entirely. The AIGE model is publicly available in python and can be accessed here: https://github.com/calvinrboyce/ai_gene_enrichment.

## 2 Results

### 2.1 Model Architecture

The AIGE model takes a list of genes as input as well as (optionally) a set of background genes to enrich against, a list of search terms for the PubMed literature search, and a brief context to provide the LLM to assist in report generation. It first sends the gene list and background set (if provided) to ToppFun[5], Enrichr[6], and g:Profiler[8] to obtain Over-Representation Analysis (ORA) results from databases including the Gene Ontology[3], KEGG[10], Reactome[27], WikiPathways[28], protein-protein interactions, and several cell atlases. Each tool uses slightly different statistical corrections to the classic hypergeometric test, with various strengths and weaknesses, so the AIGE model integrates *p*-values from all three tools to create an ensembled set of ORA results across each database. It then queries NCBI’s PubMed database to find recent articles that mention subsets of the provided genes, as well as optionally provided search terms (e.g. a paper studying glioblastoma that mentions 4 of the provided genes in an identified process). If the gene list is differentially ranked, or if the list is shorter than 10 genes, the top genes (up to 10) are queried against NCBI’s gene database to obtain summaries for each gene. ORA results from 12 databases (as obtained by the three statistical tools), literature results from PubMed (article titles, abstracts, and paragraphs), as well as gene summaries are then provided to an LLM (OpenAI’s gpt-4.1-mini-2025-04-14)[29] to identify functional themes. Highly statistically significant ORA results are aligned with related literature findings to create a list of functional themes enriched in the input list of genes, each with a description of what the theme does biologically and why it was identified. The LLM also generates a confidence score between 0 and 1 reflecting the coherence and significance of each theme. The AIGE model then produces a high level summary of all the functions found to be enriched in the input.

### 2.2 Model Evaluation

In the absence of experimental ground truth, we sought to validate the AIGE model by evaluating its ability to recreate labels for known lists of genes in several databases. Specifically, we evaluated its ability to recreate biological functions using the Biological Process branch of the Gene Ontology[3] and the Hallmark dataset from MSigDB[11], pathways using the Reactome dataset[27], and cell types using marker genes obtained from PanglaoDB[30]. We then evaluated the generated confidence scores to understand how the AIGE model assesses its own confidence by running the model on gene sets contaminated with randomly sampled genes (as in Hu et. al.[17]).

#### 2.2.1 Biological Process Identification

To validate the model’s ability to identify biological processes, we tested it on gene lists obtained from the Gene Ontology Biological Process database as well as MSigDB’s Hallmark dataset. Each gene list was provided to the model, which generated a list of themes found to be enriched in the gene set. Each of those themes was then compared to the database’s name for the gene list and evaluated semantically to determine how similar the proposed function was to the true function (following the work of Hu et. al.). We generated a percentile score for each term representing the similarity of the proposed name to the true name, as compared with all biological function names in the database. Thus, a percentile score of 95 indicates that the function proposed by the AIGE model is more similar to the database’s labelled function than it is to 95% of all biological functions represented in the database.

Because the model has access to the GO database via its ORA tools, for each GO term we first blinded the model to the specific input term, then ran the analysis with masking so it couldn’t cite the exact input term in its results. Due to the hierarchical nature of the GO database, the model still had access to parent and child terms in the GO graph, resulting in slight data leakage, so performance was expected to be high. The model was not provided search terms, background genes, or context during testing. When evaluated on 500 randomly sampled gene sets from GO with lists of genes ranging from 5 to 500 genes in length, the AIGE model was able to recover biological processes above the 95th percentile for 93% of terms (466 terms), with a median percentile score of 99.94.

When evaluating on the MSigDB database, the model was completely blinded to MSigDB as a data source to eliminate data leakage. Because the MSigDB hallmark dataset only contains 50 terms, the model’s proposed names were semantically compared to the 50 MSigDB terms as well as the 30,786 GO Biological Process term names for a more representative semantic distribution. Of the 50 gene sets, the AIGE model was able to recover biological processes above the 95th percentile for 70% of terms (35 terms), with a median percentile score of 99.85.

#### 2.2.2 Pathway Identification

Using the same validation scheme as above, we similarly evaluated the model on its ability to reproduce pathways from the Reactome database. Of the 2785 pathways in the database, we randomly sampled 480 terms to test. We blinded the model to the Reactome database as a whole (though it still had access to KEGG and WikiPathways), and computed percentile scores against the full list of pathways. The model was not provided search terms, background genes, or context during testing. The AIGE model was able to recover pathways above the 95th percentile for 86% of terms (413 terms) with a median percentile score of 99.75. In a successful case, the Reactome term “Zygotic genome activation (ZGA)” was mapped to the AIGE model’s proposed theme of “Embryonic Development and Zygotic Genome Activation (ZGA)”. In an unsuccessful case, the Reactome term “Post-translational protein phosphorylation” was mapped to “Signaling and Receptor Activity”.

#### 2.2.3 Cell Typing

Though the AIGE model was not designed to perform cell typing, it does have access to multiple cell atlases, and we evaluated its performance identifying 178 cell types using marker genes in the PanglaoDB dataset. Since the model does not have access to the PanglaoDB atlas, no masking was required during testing. The model was not provided search terms or background genes during testing, but the context phrase “Include a theme for likely cell type” was provided. Of the 178 cell types tested, the AIGE model was able to recover cell types above the 95th percentile for 77% of terms (137 terms) with a median percentile score of 99.44.

#### 2.2.4 Error Characterization

For each test database we examined the 10 worst performing terms to gain insight into the types of mistakes made by the model and identified several mechanisms for failure.

First, the semantic embedding model used to select the most likely theme from the output was at times prone to error. For example, for the GO term “Maintenance of lens transparency” (GO:0036438), the AIGE model identified several functional themes including “CHMP4B and Cellular Membrane Dynamics and Cataractogenesis”, but the semantic comparison instead selected the “Tissue and Anatomical Structure Homeostasis” theme as the most semantically similar to the GO term name. The comparison may have weighed the semantic similarity of “maintenance” and “homeostasis” more highly than the biological similarity of “lens transparency” and “cataractogenesis”, a problem which would readily be avoided when a human reviews the output of the model.

Second, small annotations sometimes led the model to pick up on more general topics than labeled in the database. For example, The GO term “Positive regulation of granulocyte macrophage colony-stimulating factor production” contains only 15 genes, for which the AIGE model proposed “Cytokine Production and Immune Response Regulation”. The model correctly identified the involvement of these genes with cytokine production but was unable to correctly identify GMCSF. These 15 genes are likely involved in the regulation of many cytokines, thus the proposed name was not incorrect but lacked specificity.

Third, in some cases the heterogeneous nature of datasets like the MSigDB Hallmark dataset led to vague or improper annotations. For example, the MSigDB term “MYC targets v1” does not accurately represent a biological function or pathway, it simply lists genes that are regulated by MYC. So the AIGE model’s proposed “Regulation of Cell Cycle and DNA Replication Initiation” theme may indeed reflect the function of the genes listed, even if it is not similar to the term name.

Fourth, database labels were at times ambiguous or inaccurate as in the case of the PanglaoDB cell type “Undefined placental cells”. In these cases, the model’s proposed “Imprinted Genes and Thyroid Hormone Metabolism - DIO3 and H19 Region” theme may be a more accurate annotation of the gene list than the assigned label.

#### 2.2.5 Confidence Evaluation

To contextualize the confidence scores provided for each theme in the model’s output, we examined the distribution of confidence scores produced across all test cases in all four databases (GO, MSigDB, Reactome, PanglaoDB), and determined distinct confidence modes (Figure 6a). Based on this distribution, we binned confidence scores above 0.93 as “High Confidence,” scores between 0.87 and 0.93 as “Medium Confidence,” and scores lower than 0.87 as “Low Confidence”. We then sampled 200 of our 480 Reactome test cases and generated a 50/50 mix test case (with half real genes from the database term and half randomly sampled genes from the superset of genes in the database), and a truly random test case (a random sample of genes from the database with the same length as the original gene list) for each test case. The AIGE model proposed high or medium confidence in 87% of pure test cases, 82% of mixed test cases, and 24% of random test cases, exhibiting only a mild reduction in confidence for the mixed cases, but a satisfactorily large reduction for random cases. However, analyzing the model’s accuracy for those same test cases revealed that the AIGE model recovered terms in the 95th percentile or higher for 86% of pure test cases, 84% of mixed test cases, and 28% of random test cases, indicating that while confidence didn’t decrease much for the mixed cases, neither did performance. At a 50% Signal to Noise Ratio (SNR), the AIGE model’s performance only dropped by 2%. The percentage of test cases labeled as high or medium confidence (87%, 82%, 24%), tracks closely with the accuracy of the model (86%, 84%, 28%), indicating that the confidence assessment is correctly evaluating performance.

**Fig. 1.**
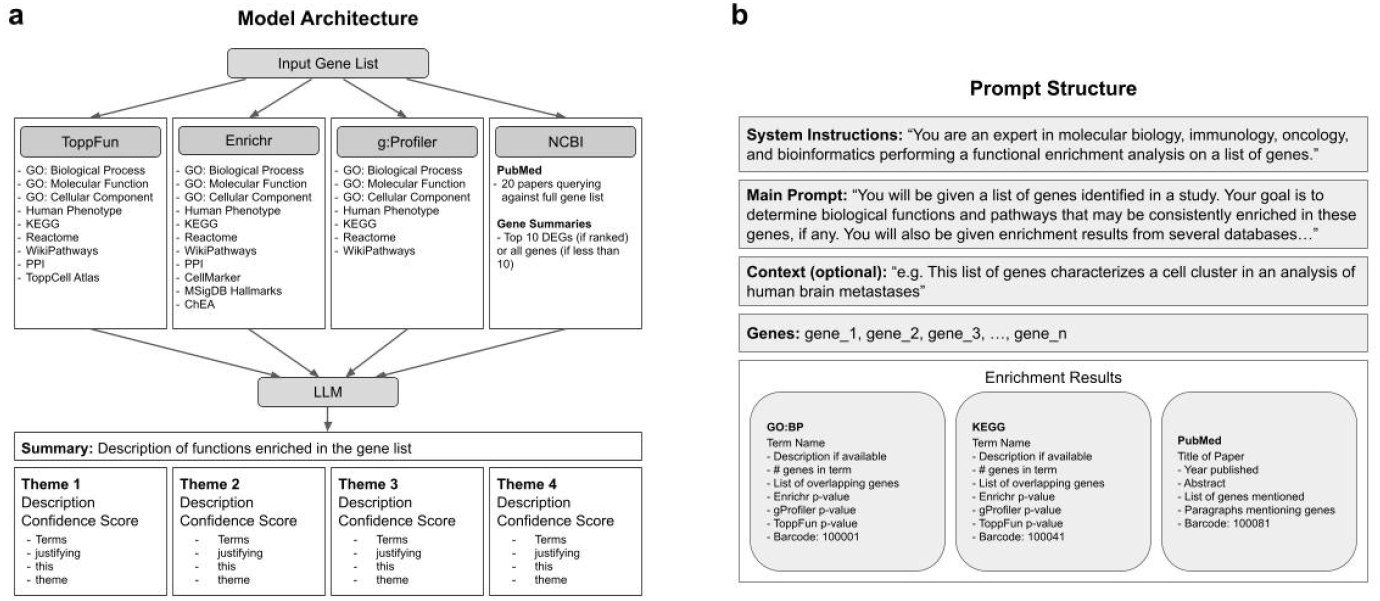
**a**, The AIGE model performs functional gene enrichment by sending the input list of genes to three statistical tools, ToppFun, Enrichr, and g:Profiler, as well as NCBI to fetch literature results, then sends all data to an LLM to identify themes in the results. **b**, The structure of the prompt includes system instructions, the main prompt, optional context, the list of genes, and the enrichment results.

**Fig. 2.**
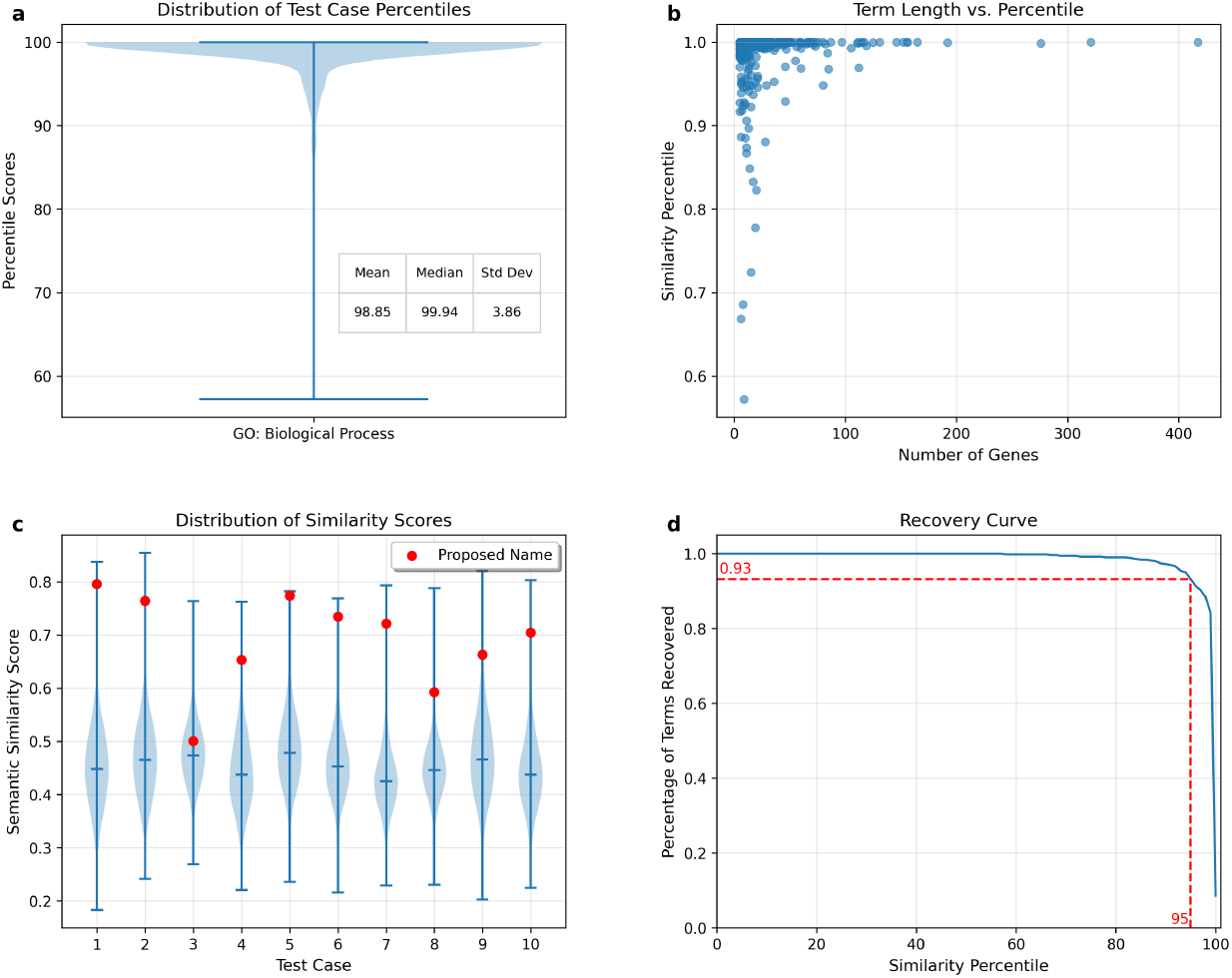
**a**, The distribution of percentile scores for names generated by the AIGE model. 500 Gene Ontology terms were tested. **b**, The relationship between the length of the input gene list and the resulting percentile score. **c**, The distributions of semantic similarities between the proposed name and each term in the database for 10 randomly sampled test cases. The correct term name is indicated with a red dot. Terms displayed are: 1) Negative regulation of signal transduction by p53 class mediator, 2) Negative regulation of axon regeneration, 3) Maintenance of lens transparency, 4) Negative regulation of smoothened signaling pathway, 5) Macrophage activation, 6) Positive regulation of activated T cell proliferation, 7) Glycoprotein biosynthetic process, 8) Tricuspid valve morphogenesis, 9) Negative regulation of fatty acid biosynthetic process, 10) Sperm flagellum assembly. **d**, The percentage of database terms recovered at a given similarity percentile threshold. The number of terms recovered above 95% similarity is indicated in red.

**Fig. 3.**
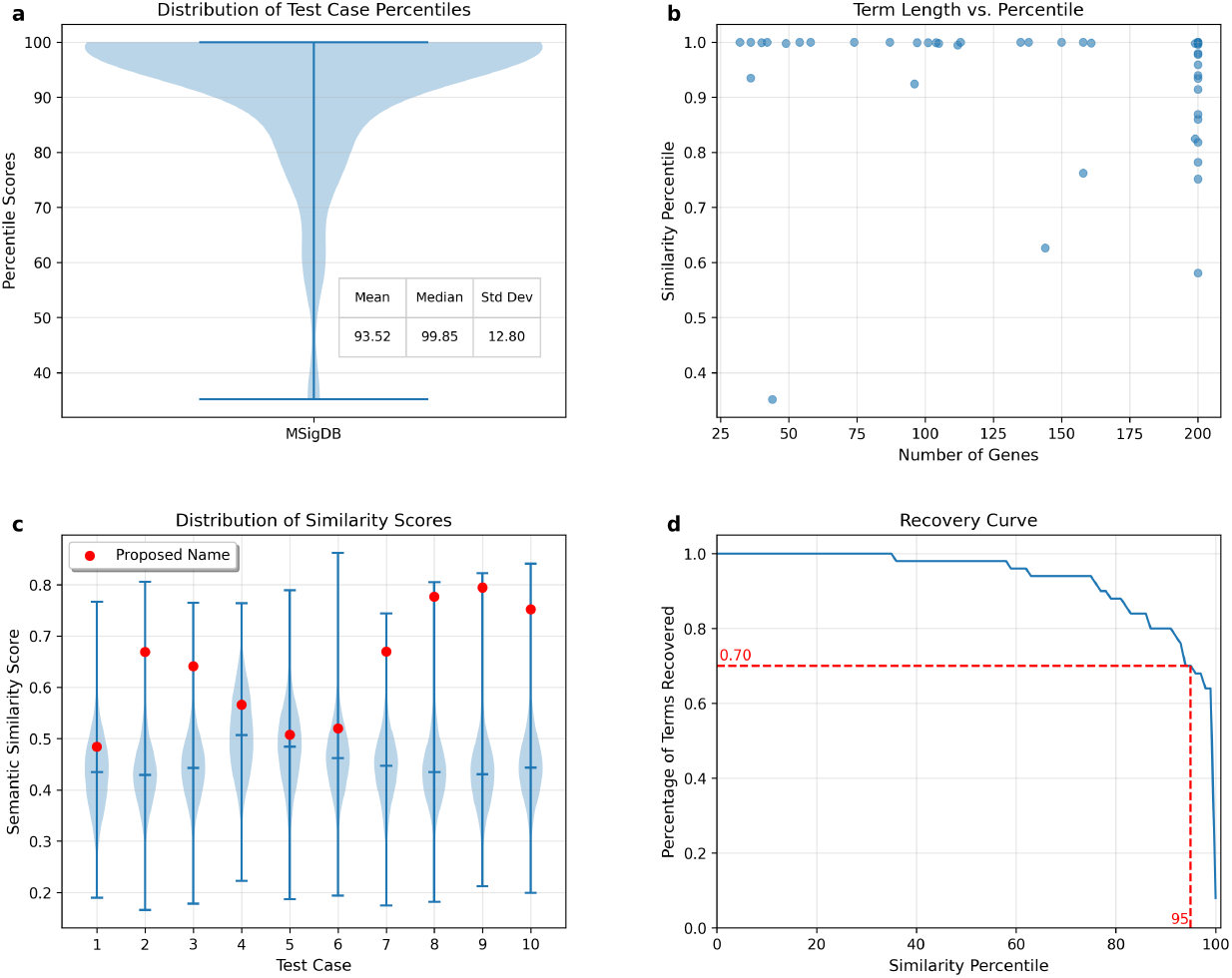
**a**, The distribution of percentile scores for names generated by the AIGE model. 50 MSigDB terms were tested. **b**, The relationship between the length of the input gene list and the resulting percentile score. **c**, The distributions of semantic similarities between the proposed name and each term in the database for 10 randomly sampled test cases. The correct term name is indicated with a red dot. Terms displayed are: 1) UV response up, 2) Bile acid metabolism, 3) Estrogen response early, 4) KRAS signaling up, 5) UV response dn, 6) Mitotic spindle, 7) Reactive oxygen species pathway, 8) Heme metabolism, 9) Notch signaling, 10) Peroxisome. **d**, The percentage of database terms recovered at a given similarity percentile threshold. The number of terms recovered above 95% similarity is indicated in red.

**Fig. 4.**
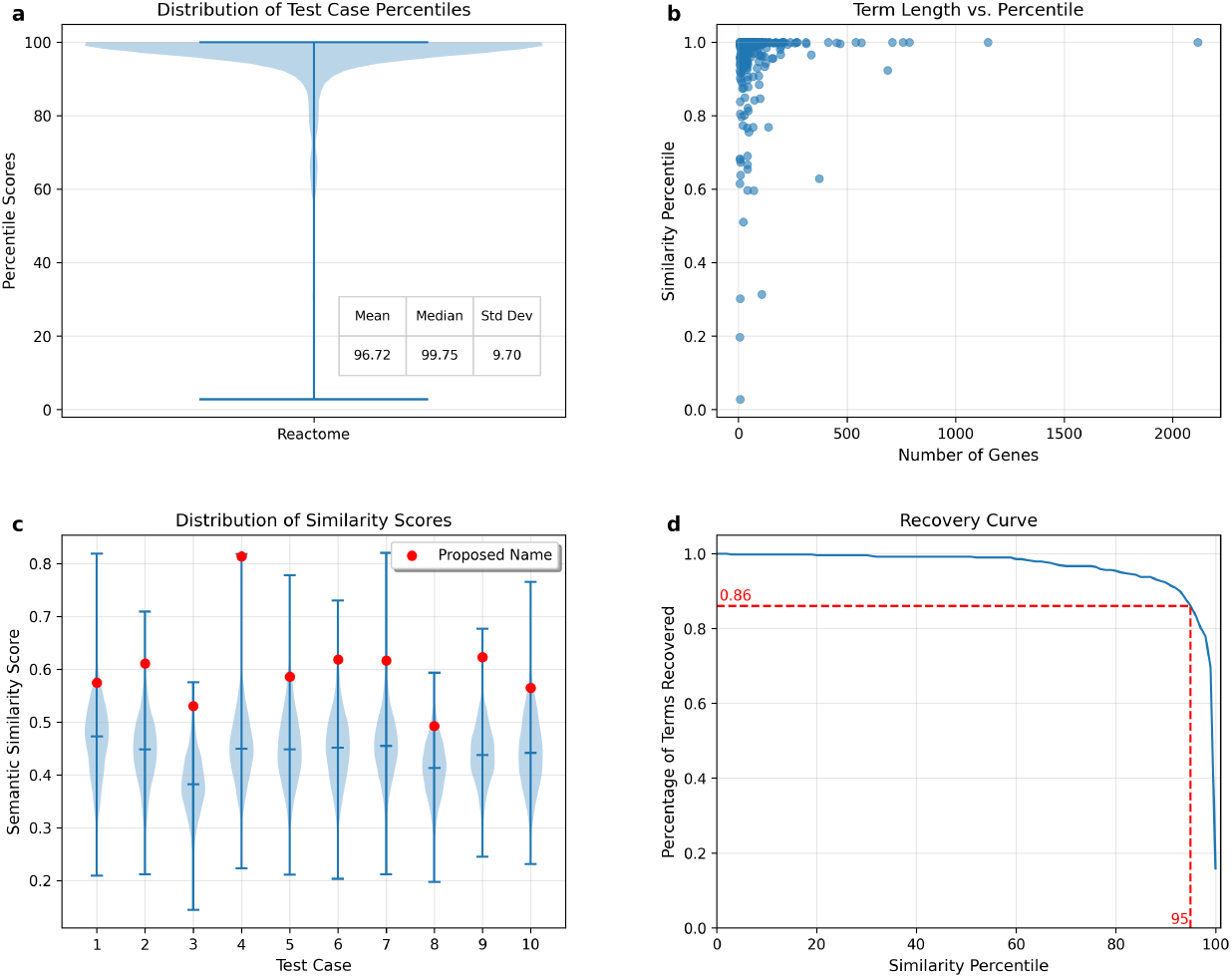
**a**, The distribution of percentile scores for names generated by the AIGE model. 480 Reactome terms were tested. The median percentile score is marked in red. **b**, The relationship between the length of the input gene list and the resulting percentile score. **c**, The distributions of semantic similarities between the proposed name and each term in the database for 10 randomly sampled test cases. The correct term name is indicated with a red dot. Terms displayed are: 1) RND1 GTPase cycle, 2) Nuclear events stimulated by ALK signaling in cancer, 3) Mtb iron assimilation by chelation, 4) Other semaphorin interactions, 5) Signaling by high-kinase activity BRAF mutants, 6) Dual incision in TC-NER, 7) Neurotransmitter release cycle, 8) Positive Regulation of CDH1 Gene Transcription, 9) Removal of aminoterminal propeptides from gamma-carboxylated proteins, 10) Fatty Acids bound to GPR40 (FFAR1) regulate insulin secretion. **d**, The percentage of database terms recovered at a given similarity percentile threshold. The number of terms recovered above 95% similarity is indicated in red.

**Fig. 5.**
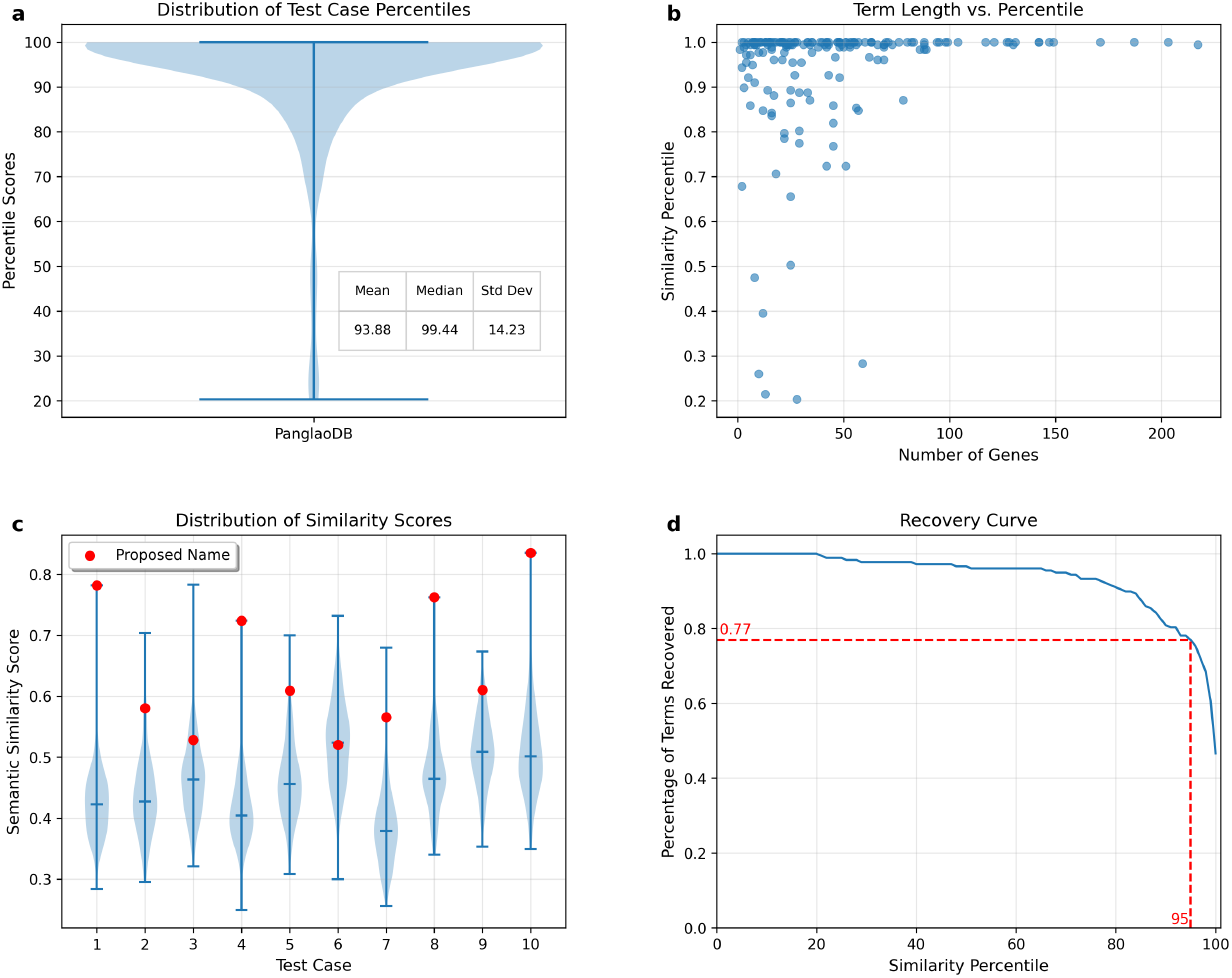
**a**, The distribution of percentile scores for cell types generated by the AIGE model. 178 cell types were tested. The median percentile score is marked in red. **b**, The relationship between the number of provided marker genes and the resulting percentile score. **c**, The distributions of semantic similarities between the proposed cell type and each cell type in the database for 10 randomly sampled test cases. The correct cell type is indicated with a red dot. Cell types displayed are: 1) NK cells, 2) Hepatic stellate cells, 3) Ependymal cells, 4) Serotonergic neurons, 5) Pericytes, 6) Parathyroid chief cells, 7) Bergmann glia, 8) Dendritic cells, 9) Microfold cells, 10) Radial glia cells. **d**, The percentage of cell types recovered at a given similarity percentile threshold. The number of cell types recovered above 95% similarity is indicated in red.

**Fig. 6.**
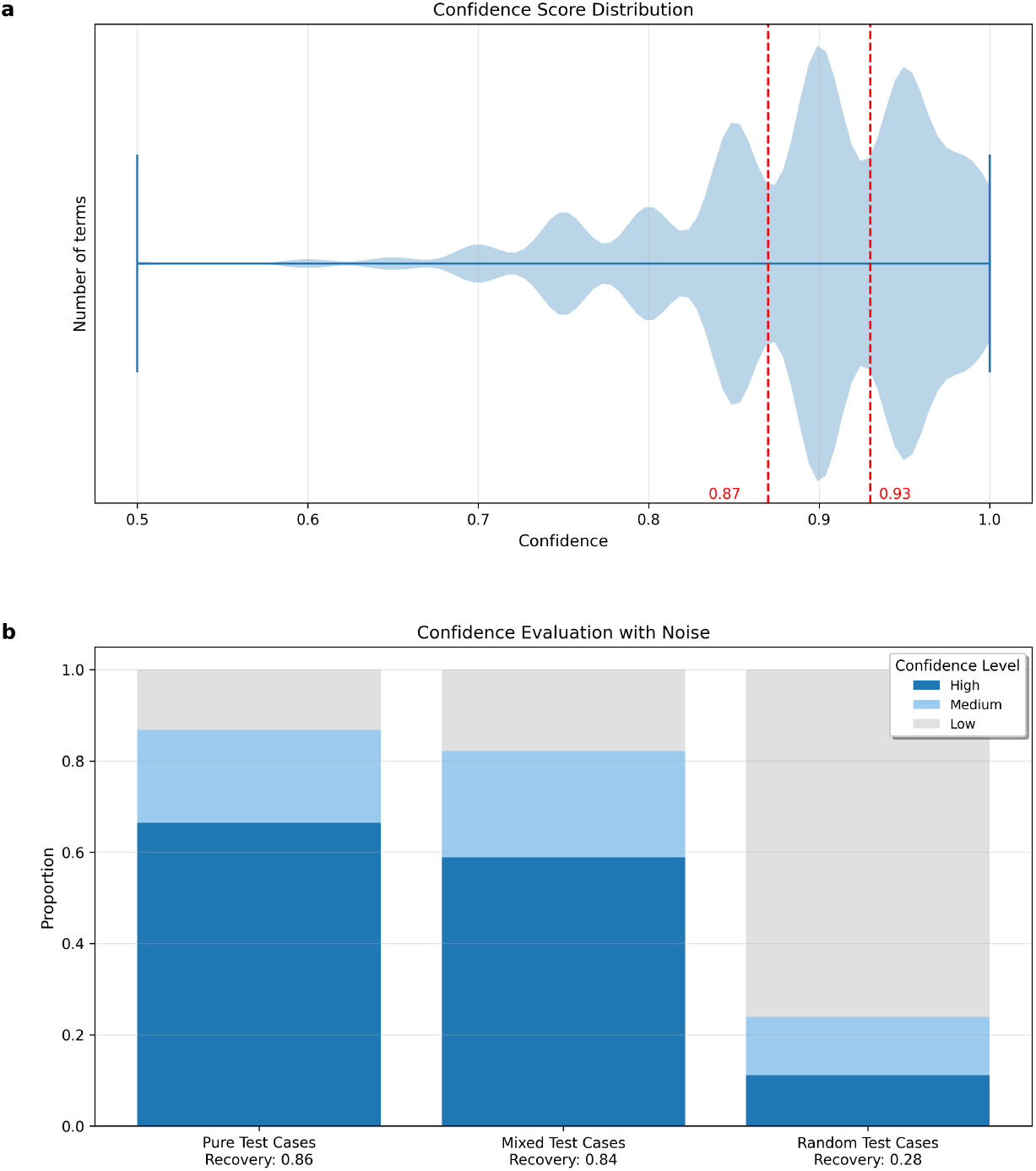
**a**, The distribution of confidence scores provided across all four test datasets, Gene Ontology, MSigDB, Reactome, and PanglaoDB. Distinct confidence modes were identified and segmented into “High,” “Medium,” and “Low” confidence with dotted red lines. **b**, Percentage of test cases assigned “High,” “Medium,” and “Low” confidence for the pure tests (100% real genes), mixed tests (50/50 real and random), and random tests (100% random genes).

## 3 Discussion

Functional gene annotation requires careful examination and subjective interpretation of a large amount of data from statistical and literature sources, a task that can naturally be extended with LLMs. The AIGE model’s workflow carefully reflects the best practices of human experts to curate an enrichment report that accurately reflects the biological functions and pathways enriched in novel gene lists. The model was able to accurately recover 93% of biological functions, 86% of pathways, and 77% of cell types across a total of 1208 test cases. AIGE was also able to accurately assess its own self-confidence with high and medium confidence percentages accurately reflecting term recovery. Analysis of model confidence also revealed a robust capacity for AIGE to correctly label biological pathways in the presense of noisy data.

The AIGE model is able to quickly parse textual data from a greater number of databases than a human would reasonably be able to analyze. However, it is still limited to the terms labeled in the current databases and findings published in literature. It is unable to generalize to novel functional annotations not previously documented. It is also subject to any errors present in the curation of these datasets, including poor labeling and incomplete gene lists. The volume of data consumed by the model tends to outweigh these insufficiencies, but it does not solve the problem entirely.

Functional gene annotations via tools such as ToppFun, Enrichr, and g:Profiler are also subject to errors induced by the statistical assumptions of Over-Representation Analysis. Each of the three tools used for statistical comparison still assumes a uniform distribution of human genes and an independent assortment of those genes, both of which are evidently incorrect assumptions. Each tool attempts to account for these assumptions with a unique statistical correction, an ensemble of which provides AIGE a more robust representation of functional enrichment. However, a more intricate statistical comparison would provide more accurate enrichment results.

## 4 Methods

### 4.1 ToppFun

ToppFun is part of the ToppGene Suite, and performs ORA via a hypergeometric test with Bonferroni or FDR correction[5]. We used their published Gene Symbol Lookup API[31] to convert gene symbols to their Entrez IDs, then their Functional Enrichment API[32] to retrieve ORA results. The API does not currently support background gene lists, so we were unable to programmatically enrich against a background gene list. The API returned data for category (GO:BP, GO:MF, GO:CC, HumanPheno, etc), ID, name, p value, various statistical corrections to p values, data on number of genes in the provided list as well as the database term list and their overlap, and a list of genes found in both lists. We included results from the GO: Biological Process, GO: Molecular Function, GO: Cellular Component, Human Phenotype, KEGG, Reactome, WikiPathways, Protein Protein Interaction, and ToppCell Atlas databases.

### 4.2 Enrichr

Enrichr performs ORA via the Fisher exact test, and then corrects the ranking results by comparing them against a standard distribution of rankings to see how each term deviates from its expected rank[6]. We used their published addList API to add the input gene list to their database, the addbackground API to upload a background gene list to enrich against, and their enrich API to retrieve ORA results[33]. The API returned data for rank, term name, p-value, several statistical corrections to p values, and overlapping genes. We included results from the GO: Biological Process, GO: Molecular Function, GO: Cellular Component, Human Phenotype, KEGG, Reactome, WikiPathways, Protein Protein Interaction, CellMarker, MSigDB Hallmarks, and ChEA databases.

### 4.3 g:Profiler

g:Profiler performs ORA via a hypergeometric test and corrects for multiple comparisons using their g:SCS (Sets, Counts and Sizes) method[8]. We used their python package to send gene list queries as well as background gene sets and compute ORA results[34]. The python package returned data for term name, ID, description, p-value, and query and term size and overlap. We included results from the GO: Biological Process, GO: Molecular Function, GO: Cellular Component, Human Phenotype, KEGG, Reactome, and WikiPathways databases.

### 4.4 NCBI

The literature search was performed via NCBI’s Entrez ESearch[35] and EFetch[36] utilities to find PubMed papers that mention individual genes or subsets of genes from the input list. The utilities were programmatically accessed via the Biopython Entrez package[37]. We queried the entire list of genes against the PubMed database to find the top 20 papers that mention genes from the input list, as well as an optional list of provided search terms. Queries were generated to find papers that mention one or more genes in the Text Words [tw] and one or more of the optional search terms in the Medical Subject Headings [MeSH] if provided, and that were published since 2015 as follows:

~~~
“(gene_1[tw] OR gene_2[tw] OR gene_3[tw]) AND (term_1[MeSH Terms] OR term_2[MeSH Terms]) AND 2015:3000[PDAT]”
~~~

The names, publishing dates, and abstracts of each paper were stored, as well as individual paragraphs mentioning genes from the input if the paper was available on PubMed Central for public access. All text was scraped for mentions of the genes, which were then surrounded by asterisks to bold the gene mentions for the LLM (`gene_1` became ^**^gene_1^**^). Genes mentioned in the text were also stored in a list of mentioned genes.

For ranked lists, the ESearch and ESummary utilities were used to obtain gene summaries of each of the top 10 genes. For gene lists less than or equal to 10 genes, all genes were queried for gene summaries.

### 4.5 Data Presentation

The top 20 results from each database (ranked by lowest *p*-value returned by any enrichment tool) were passed into the LLM for synthesis, along with all PubMed results. Enrichment data were presented to the LLM in JSON format including lists of dictionaries for each database. For the ORA databases, each term’s dictionary contained the term name, description, overlapping genes, term size, and p-values from each of the applicable tools (ToppFun, Enrichr, g:Profiler), as well as a uniquely generated barcode for the LLM to identify the term. PubMed papers included the name, publishing year, abstract, list of mentioned genes, and quotes of specific gene mentions if publicly available, as well as a uniquely generated barcode.

#### GO:BP

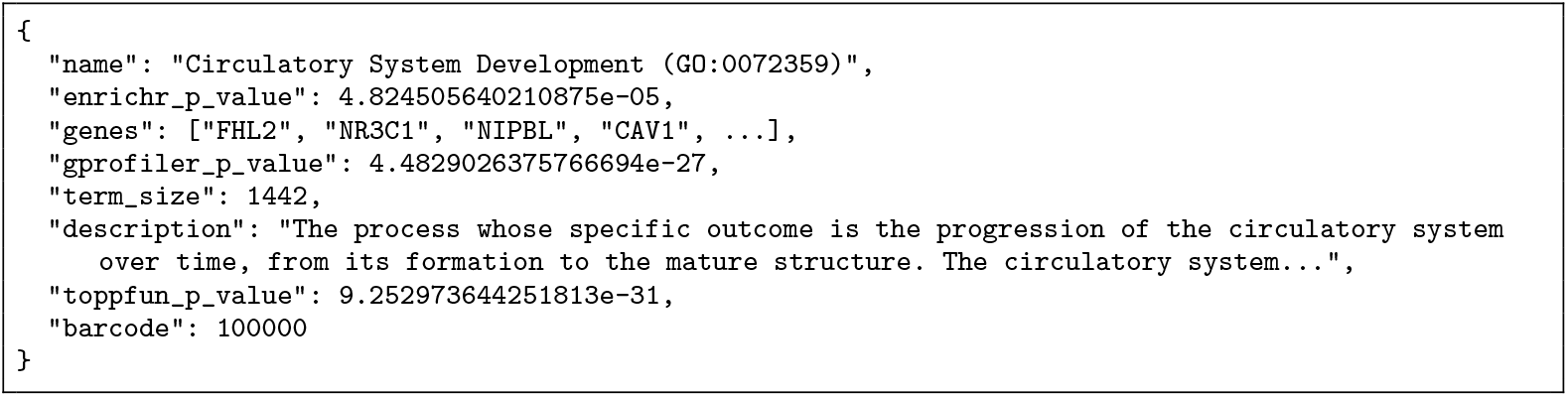

#### PubMed

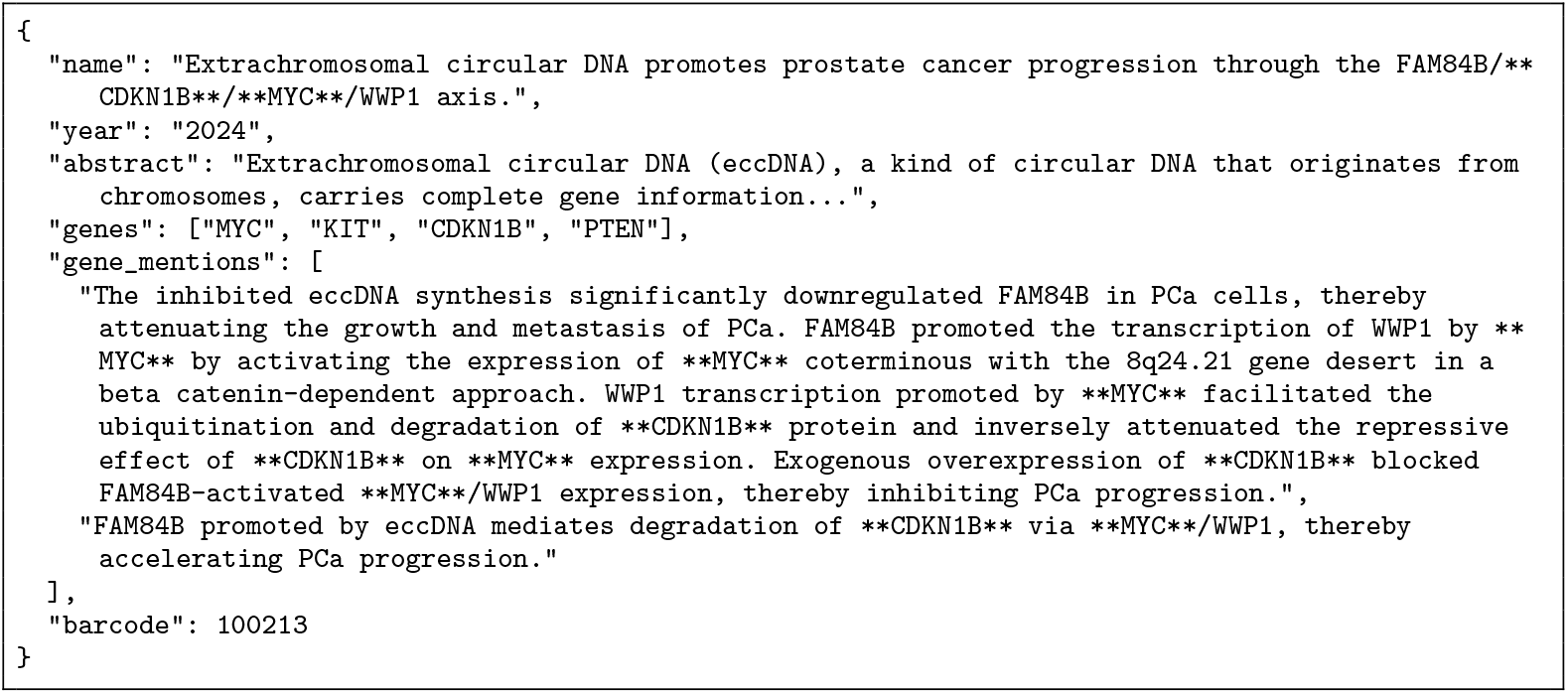

### 4.6 LLM Prompting and Output Parsing

Instructions were given to the LLM in 5 parts as follows. (1) System Prompt: The model was instructed to behave as an expert in molecular biology, immunology, oncology, and bioinformatics. This section of the prompt informs the personality, style, and specificity of the output. (2) Main Prompt: Here the model was informed of its task to determine biological functions and pathways enriched in the provided set of genes. It was provided context on the sources of enrichment results it would be parsing and instructed to organize them into functional themes that accurately represent the functions enriched in the genes. It was also instructed to output a list of themes, each with a theme name and description as well as a confidence score based on the p-values of the enrichment terms and the number of genes involved in the theme. Due to the tendency of LLMs to hallucinate things like term IDs and p-values, instead of asking the model to reproduce the enrichment terms related to the theme, it was simply asked to list the integer barcodes of the terms that support each theme. This eliminates the potential for the model to hallucinate information that wasn’t provided in the initial search. (3) Context: Input to the model can optionally include brief context to help it focus on enrichment results relevant to the current study. For example, “This list of genes characterizes a cell cluster in an analysis of human brain metastases”. (4) Genes: The input genes were then listed separated by commas. (5) Enrichment Results: Results from the various databases were dumped in JSON format as described above.

The above LLM prompt was provided to the “gpt-4.1-mini-2025-04-14” model via OpenAI’s Responses API[38] at a price of $0.40 per 1M input tokens and $1.60 per 1M output tokens, averaging $0.022 per run. To minimize hallucination and ensure consistent data parsing, the LLM was provided with a pydantic model as a Structured Output[39] to retrieve data according to a proper JSON schema:

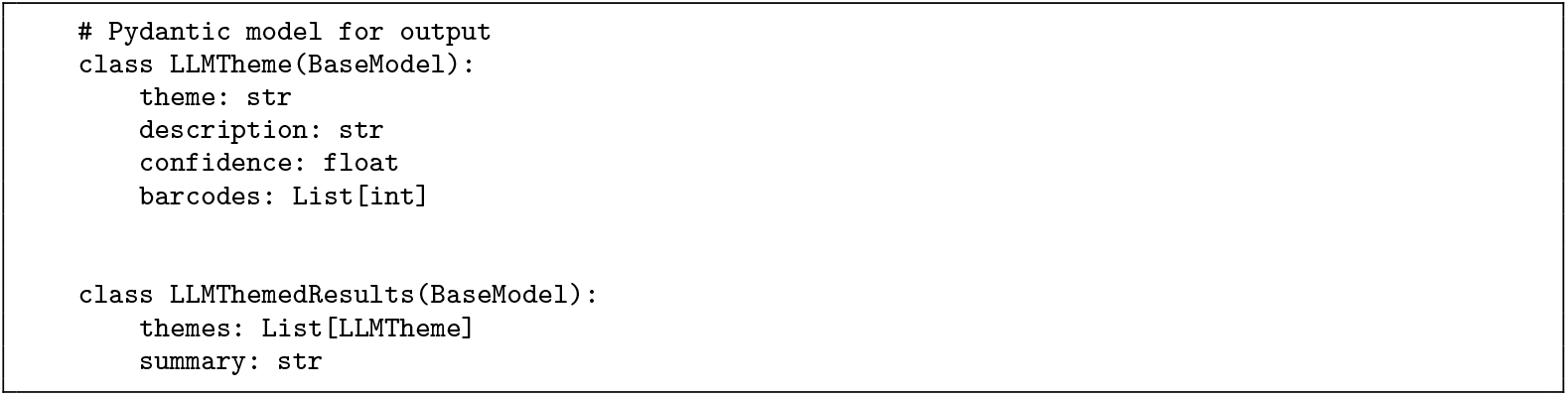

The listed barcodes were then checked against the original enrichment results to fetch the IDs and p-values to eliminate data hallucination.

### 4.7 Formatted Presentation

For readability, results are optionally written into an Excel spreadsheet with a summary page that includes metadata from the run as well as the overall summary and list of themes identified, followed by a page for each identified theme that includes the description of the theme, the confidence score, and the individual enrichment results that support the theme with their IDs and p-values.

### 4.8 Gene Set Validation Scheme

To assess the ability of our model to extract accurate functional annotations from gene lists, we validated its performance on curated databases of gene lists and their known functions using the same validation framework as found in Hu et. al.[17] We obtained data from the Gene Ontology: Biological Process[40, 41], MSigDB Hallmarks[42], Reactome Pathways Gene Set[43], and PanglaoDB Marker Genes[44] data sets. Each database consists of a set of functions *Y* = {*y*_*i*_}, where each *y*_*i*_ is a label, and a set of gene lists *S* = {*X*_*i*_}, where each *X*_*i*_ = {*x*_*j*_} is a set of genes known to be involved in the function labeled by *y*_*i*_. For a single term *y*_*i*_, after running ORA and a literature search on the corresponding genes *X*_*i*_, the model proposed a list of 5-10 themes identified in the data, *Ŷ*_*i*_ = {*ŷ*_*j*_ } As identified in the work of Hu et. al., each list of genes may be properly enriched for multiple biological functions besides the one labeled in the database, so we first semantically compared each proposed theme *ŷ*_*j*_ ∈ *Ŷ*_*i*_ with the true term label *y*_*i*_ to find the proposed theme *ŷ*_*i*_ most likely to represent the true label *y*_*i*_. The proposed label *ŷ*_*i*_ was then semantically compared to all terms in the source database *Y*, to create a similarity distribution representing how closely the proposed label *ŷ*_*i*_ reflected each biological function in the database. We then computed a percentile for the true label *y*_*i*_. Thus a percentile score of 95% means that the proposed function represents the true function more closely 14 than 95% of all biological functions in the database. As in Hu et. al., all semantic comparisons were performed by cosine similarity of SapBERT[45] transformer embeddings, which were computed via the transformers package from Huggingface[46] (cambridgeltl/SapBERT-from-PubMedBERT-fulltext).

## Code Availability

Code for the AIGE model was written in python and partially generated with Cursor[47], an LLM-augmented IDE. All code used for testing has been made available at https://github.com/calvinrboyce/gene enrichment agent. A production version of the AIGE model is available at https://github.com/calvinrboyce/ ai gene enrichment.

## Notes

### Competing Interest Statement

The authors have declared no competing interest.

### Summary of Updates

Main updated to address recent literature, code documentation updated

### Funding

- Simmons Center for Cancer Research

